# Examining the Central Auditory Processing Using a Multi-Feature ERP Paradigm to Phonetic, Prosodic and Acoustic Features

**DOI:** 10.1101/2024.07.23.604852

**Authors:** Zavogianni Maria Ioanna, Maja Kelić, Ferenc Honbolygó

## Abstract

Previous studies have shown that multi-feature paradigms can be used for the investigation of the central auditory processing. In this study, we developed a speech multi-feature paradigm with phonetic, prosodic, and acoustic changes. Our aim was to examine the participants’ involuntary discrimination of the changes of speech sound features while they were watching a movie without any sound or subtitles. All five deviant conditions (vowel, consonant, stress, intensity, frequency) elicited statistically significant MMN ERP responses which varied in amplitude depending on the condition; stress was also connected with the occurrence of the LDN component. With vowel, consonant and stress among the conditions that received the most statistically significant ERPs, we could suggest that the current multi-feature paradigm can be used to successfully elicit the MMN ERP component and potentially to assess phonological processing in children and adults.

## Introduction

Event-Related brain Potentials (ERPs) is a time-sensitive Electroencephalography (EEG) technique which is used to study language processing (Kaan, 2007). One of the ERP components, the Mismatch Negativity (MMN) ERP component, has been continually used for the investigation of speech perception (Luck, 2014). Being elicited within an auditory condition, the MMN component is considered as the brain’s electrical response to auditory alterations since it is evoked by a discriminable change (i.e., a ‘deviant’ stimulus) during the presence of regular auditory features (i.e., a ‘standard’ stimulus) (Näätänen et al., 2007). The MMN component is characterized by negative polarity and fronto-central brain activity distribution, and occurs in a time window of approximately 150 ms to 250 ms post the deviant onset (Luck, 2014).

Typically, the experimental design which allows the recording of MMN responses is the passive oddball paradigm, in which the standard stimulus occurs with a high probability (cca. 80%) and a single deviant stimulus appears with a low probability (cca. 20%). While this is a reliable method to elicit the MMN component, it is very time consuming, and not ideal to use with special (i.e., infants, typically developed children, children with reading disorder) or clinical populations (i.e., patients with Parkinson’s and Alzheimer’s disease). In order to tackle this issue, a more optimal version of the paradigm has been suggested by Näätänen et al. (2004), the so-called multi-feature paradigm. In this paradigm, one standard stimulus and several deviant stimuli are presented in an auditory manner. Pakarinen and his collaborators conducted both a passive oddball and a multi-feature paradigm in order to compare them with one another (Pakarinen et al., 2009) and reported that the MMN peaks recorded during the multi-feature paradigm were equally reproducible with the MMN peaks recorded during the passive oddball paradigm (Pakarinen et al., 2009). During the speech multi-feature paradigm, the MMN ERP response is recorded with the participants’ involuntary activity (i.e., the participants can engage in activities, such as watching a film without sound). This occurs since the MMN ERP responses are generated via an automatic brain process, thus they are not attention-dependent (Näätänen, Paavilainen, Rinne & Alho, 2007).

Previous studies have applied the multi-feature paradigm in order to investigate the MMN ERP responses evoked by alterations on the phonetic or acoustic parameters of the syllables of the certain stimuli (Pakarinen et al., 2009; Pakarinen et al., 2013; Pakarinen 2014; Sorokin, Alku & Kujala, 2010; Partanen, Vainio, Kujala, and Huotilainen, 2011; Fisher et al., 2011; Kuuluvainen et al., 2014; David et al., 2020). The new generation of multi-feature studies examining speech began with Pakarinen et al. (2009), in which the authors investigated the MMN reproducibility of the multi-feature paradigm in comparison with the oddball one. For this reason, they designed a multi-feature paradigm in which single syllabic stimuli with deviations in vowel, duration of vowel, consonant, frequency, and intensity from the standard stimulus, were used. As mentioned above, Pakarinen et al. (2009) reported that the five deviant conditions in the multi-feature paradigm elicited statistically significant MMN peaks, which were similar to the ones having been elicited during the oddball paradigm. Similarly, Sorokin et al., (2010) investigated the discrimination of vowel, consonant, syllable intensity and frequency changes, which produced statistically significant ERP responses.

To investigate whether participants were able to discriminate changes in auditory stimuli with accuracy, Partanen et al. (2011) designed an extensive linguistic multi-feature MMN paradigm with trisyllabic stimuli. Specifically, in this paradigm, the deviant conditions included changes in the duration, intensity, and frequency of the stimuli as well as a vowel change. These changes appeared in the initial, middle or the final syllable of each stimulus (Partanen et al., 2011). Even though all deviant conditions elicited significant MMN responses, the duration condition elicited the largest MMN amplitude in the changes made in the initial syllable of the stimuli because of the Finnish speakers’ sensitivity to detect stress patterns in the initial position of a word, according to the Finnish phonology (Partanen et al., 2011). Furthermore, Fisher et al. (2011), created an ‘optimal’ version (i.e., ‘Optimal – 5) of the multi-feature paradigm comparing it to a shorter version (i.e., ‘Optimal – 3). During both of the paradigms, significant MMN ERP peaks were evoked; however, the deviant conditions of the optimal multi-feature paradigm elicited the largest MMN amplitudes confirming that a multi-feature paradigm with 5 deviant stimuli increases the probability of larger MMN responses (Fisher et al., 2011).

Furthermore, Pakarinen et al. (2013) evaluated the central speech-sound processing with MMN ERP measurements. They designed a multi-feature paradigm in which the deviant conditions included changes in vowel duration, intensity level, and frequency of the stimuli as well as vowel changes (Pakarinen et al., 2013). Even though the deviant conditions elicited statistically significant MMN ERP responses, differences in the peaks were reported according to the vowel duration (i.e., short or long categories of a phoneme). Pakarinen et al. (2013) suggested that this multi-feature paradigm can be used to study phoneme categorization especially for languages which are dependent on phonemic variations (e.g., Finnish). In 2014, Pakarinen and his colleagues designed a new multi-feature paradigm to examine the central speech-sound representations and the participants’ ability to pay attention to stimuli containing emotional information (Pakarinen et al., 2014). Statistically significant MMN ERP responses were reported upon the auditory presentation of all deviant stimuli (i.e., frequency, intensity, spectral density, noise level, sound-source location, consonant duration, vowel change, vowel duration, and omission of the second syllable of the standard stimulus). Additionally, large MMN peaks were found for the deviant conditions with emotional connotation (i.e., sad, happy, angry utterances) since they deviated more from the standard stimuli (Pakarinen et al., 2014). Moreover, Kuuluvainen et al. (2014) compared CV-syllable structured stimuli to acoustically representative nonspeech stimuli with respect to the occurrence of MMN/MMNm (i.e., the magnetic counterpart of MMN). They showed that the MMN/MMNm peaks were larger for the same features of both speech and nonspeech stimuli (Kuuluvainen et al., 2014); this finding provides evidence of the existence of phonetic features during the preattentive stage of cortical processing (Kuuluvainen et al., 2014).

Finally, in recent years, David et al. (2020) examined the participants’ brain responses to deviant stimuli which varied in phonological complexity (i.e., stimuli with formatted structure, such as CVCV, CCV.CV, CVC.CV). According to the results, the MMN ERP responses were of similar latency for either the simple or the complex deviant stimuli; the MMN peaks for all stimuli conditions were also statistically significant (David et al., 2020).

One of the main advantages of recording MMN ERP responses is the recording process itself since MMN is recorded in involuntary conditions and the participants participate passively (Partanen et al., 2011; Pakarinen et al., 2013). For this reason, the multi-feature paradigm has been applied to populations who are unable to participate in EEG experiments that require more traditional sound discrimination settings (Partanen et al., 2011; Pakarinen et al. 2013). Having been applied to such populations, the MMN ERP component has led researchers to study the pre-attentive cognitive responses even more thoroughly (Fitzgerald & Todd, 2020). For instance, the MMN ERP component has been extensively used in studies with patients with neurodegenerative diseases (e.g., Parkinson’s and Alzheimer’s disease; Pekkonen, 2000; Brønnick et al., 2010) or neuropsychiatric disorders, such as schizophrenia (Todd et al., 2014; Todd et al., 2018; Näätänen, Kujala & Light, 2019).

What is more, the MMN multi-feature paradigm has also been used to study language development and linguistic knowledge in infants and young children (e.g., Partanen, Pakarinen, Kujala & Huotilainen, 2013; Linnavalli, Putkinen, Huotilainen & Tervaniemi, 2018;) and monitor language performance in participants with at risk for developing language and reading disorder (Lovio, Näätänen & Kujala, 2010).

In the present study, we aimed to develop a multi-feature MMN paradigm that could be potentially used to profile the speech sound processing abilities of children with speech and language disorders. Apart from the phonetic features used in the previous studies, we also included a prosodic feature, word stress. Previously, the passive oddball MMN paradigm has been applied to study the neural processing of word stress (Weber et al., 2004; Honbolygó et al., 2020). Weber, Hahne, Friedrich, and Friederici (2004) reported that adult German speakers were able to show a typical MMN to the stress change on the first syllable of the stimulus (i.e., trochaic item) and a MMN to the stress change on the second syllable of the stimulus (i.e., iambic item). This shows that both stress patterns can be recognised.

Furthermore, a study on Hungarian (Honbolygó et al., 2004) showed that after the auditory presentation of a stimulus with stress on the second syllable, two consecutive MMN components were elicited. According to the Hungarian prosodic rules (Honbolygó, Kóbor, Borbála, & Csépe, 2020), this is an illegal stress pattern (i.e., stress on the first syllable is the legal pattern in Hungarian).

Ylinen, Strelnikov, Huotilainen, and Näätänen (2009) investigated the processing of word stress in Finnish familiar (i.e., stress on the first syllable) and unfamiliar (i.e., stress on the second syllable) stress patterns. In this study, they reported the emergence of two consecutive MMN components after the stimulus with the unfamiliar stress pattern. More specifically, this characteristic double MMN peak was later characterized as Late Discriminative Negativity (LDN) ERP component. LDN occurs after detection of changes in prosodic stimuli, and it peaks after the MMN ERP component, around 350 – 500ms. This has been frequently reported by studies on Hungarian language (Honbolygó & Csépe, 2013; Honbolygó & Csépe, 2014); for instance, Honbolygó and Csépe (2013) reported the LDN emergence after irregular stress patterns. Apart from such studies, the LDN ERP component has also occurred in the study conducted by David et al. (2020) in which the participants were native French speakers. David et al. (2020) proposed that the emergence of LDN could be a useful index of phonological complexity processing since the MMN component reflects the sound change processing at a sensory level, whereas the LDN component reflects a sound change processing at a higher cognitive level (David et al., 2020; Winkler, 2007; Ceponiene et al., 2004).

In the present study, we used the speech multi-feature paradigm to profile the speech processing abilities of participants by examining the MMN ERP component elicited by the involuntary detection of speech sound features. The paradigm was constructed with one standard and five deviant stimuli conditions including phonetic (i.e., vowel, consonant), prosodic and acoustic changes (i.e., frequency and intensity). Our aim was to study the processing of speech sound features within a multi-feature paradigm and examine how the paradigm could potentially be applied in future studies aimed at investigating special populations with speech and language deficits. More specifically, we focused on whether and how the different conditions (i.e., vowel, consonant, stress, frequency, and intensity) elicit the MMN component, and how these conditions would differ from each other with respect to the MMN activity they elicit.

This is the first study in which a multi-feature paradigm with a prosodic change as one of the deviant conditions is applied. In addition to this, it is the first study of its kind to be conducted in Hungarian. This is important because the present results add to existing findings by studying another language previously not investigated, and thus broadening the generalizability of MMN results. Moreover, the addition of prosodic features allows us to investigate a broader range of speech features, which might be an important contribution to the studies of speech and language disorders (e.g., research on prosody: Honbolygó, Csépe & Rago (2004); Ylinen, Strelnikov, Huotilainen & Näätänen (2009) and research on dyslexia: Vellutino, Fletcher, Snowling & Scanlon (2004); Pennington (2006); Snowling, Hulme & Nation (2020)).

Based on previous findings, we expected that participants would be able to involuntarily detect these phonetic and acoustic changes; in addition to this, we expected differences in the MMN activity (i.e., differences in the mean amplitudes) in each deviant condition.

## Methods

### Participants

Twenty-five participants (14 females) were recruited for the purpose of the study. The participants’ age was between 19 and 38 years (Mage =25, SD ± 5.60). All of them were native speakers of Hungarian and university students. None of them reported having any neurological disorders, hearing or speech-related issues. Prior to the start of the experiment, participants gave their written informed consent. The study was approved by the United Ethical Review Committee for Research in Psychology in Hungary (Ref. no. 2020-36).

### Stimuli

The stimuli consisted of disyllabic pseudowords in a CVCV format; the Hungarian phonotactic rules were followed for their construction (see Table 1). The standard stimulus was a pseudoword (‘keke’). The deviant stimuli were formulated by modifying the first syllable at the standard stimulus with the following phonemic and acoustic alterations.

**Table 1.**
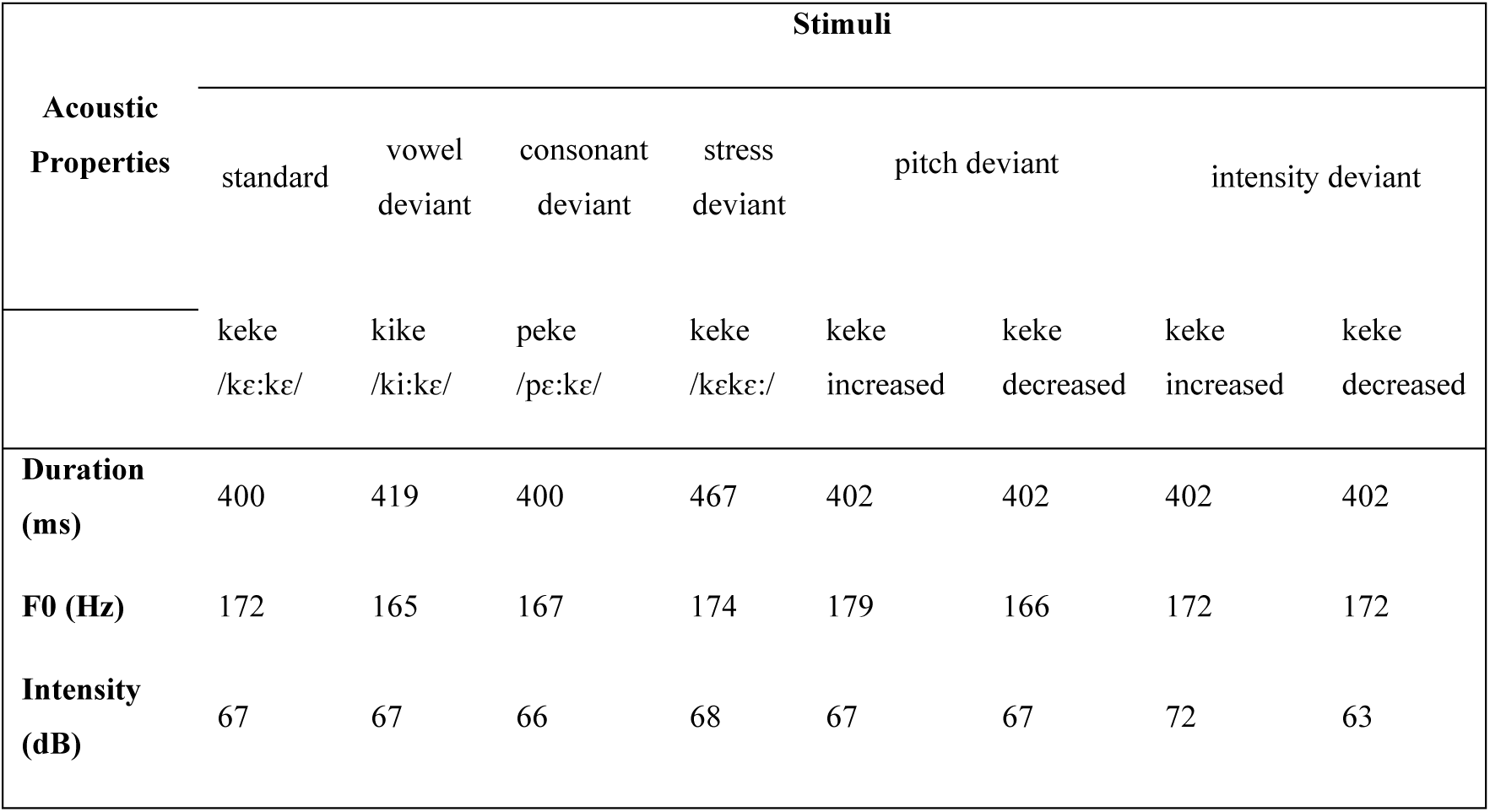
The design of the stimuli (i.e., standard, vowel, consonant, stress, pitch and intensity) together with the acoustic properties.

Specifically, the deviants consisted of a vowel deviant type (‘kike’), a consonant deviant type (‘peke’) and a stress deviant type, where the stress was on the second syllable of the stimulus (‘keke’). In addition, the deviant stimuli consisted of a frequency (F0 ± 8%) and an intensity deviant type (± 6 dB). All stimuli were recorded in isolation by a female native speaker of Hungarian in an acoustically shielded experiment booth. The stress stimulus was inserted into naturally produced utterances; after recording, the stress stimuli were isolated from the produced utterances. Finally, the stimuli were edited according to the aforementioned alterations by using PRAAT (version 6.1.52) and Audacity software.

### Procedure

The stimuli were presented in a speech multi-feature paradigm (following the multi-feature paradigm of Partanen et al., 2009). We presented a total of 615 stimuli, of which 315 were standard and 300 were deviant. The stimulus sequence began with the auditory presentation of fifteen standard stimuli; after that, each deviant stimulus was presented in a random order and in alternation with the standard stimulus. In addition, the stimuli were presented with a stimulus onset asynchrony (SOA) of 750 ms which was arranged to be randomised across the paradigm. During the EEG experiment, participants were comfortably seated in a chair inside a soundproof experiment booth; they were instructed to listen to the stimulus sequence without them being attentive to it. For this reason, participants were asked to watch a movie of their preference without any sound or subtitles while the stimulus sequence was presented to them through headphones (Sennheiser PX200). The duration of the EEG recording time was about 17 min, and the whole experiment lasted for about 90 minutes, including the application and removal of electrode caps. The presentation of the experiment was performed by using the Presentation software (v. 21.1, Neurobehavioral Systems).

### EEG Recording and Preprocessing

EEG activity was recorded with a 64-channel recording system (BrainAmp amplifier and BrainVision Recorder software, BrainProducts GmbH) with a sampling rate of 1000 Hz. The Ag/AgCI sintered ring electrodes were mounted in an electrode cap (actiCAP) on the scalp of each participant according to the 10% equidistant system at the following positions: Fp1, GND, Fp2, AF7, AF3, AFz, AF4, AF8, F7, F3, F1, F2, Fz, F4, F6, F8, FC5, FC1, FC3, FCz, FC2, FC4, FC6, FT7, FT9, FT8, FT10, T7, C5, C3, C1, Cz, C2, C4, C6, T8, TP9, TP7, CP5, CP3, CP1, CPz, CP2, CP4, CP6, TP8, TP10, P7, P5, P3, P1, Pz, P2, P4, P6, P8, PO7, PO3, POz, PO4, PO8, O1, Oz, O2, Iz. The reference and ground electrodes were mounted at the FCz and GND position, respectively. During the recording, electrode contact impedances were kept below 10 kΩ.

The EEG data was analyzed offline with BrainVision Analyzer 2.2 software (Brain Products GmbH, Munich, Germany). We started the preprocessing of the raw EEG data by applying a bandpass filter from 1 – 40 Hz (48dB/oct); in addition to this, we used the notch filter with a selected frequency of 50 Hz to remove additional electrical noise. A second step was to clean the EEG data from the horizontal and vertical (blinks) eye-movement artifacts as well as heartbeats by applying Independent Component Analysis (ICA) (Delorme, Sejnowski & Makeig, 2007); in this step, the EEG signals were split into 32 independent components, and components matching the waveform and amplitude distribution of blinks, horizontal eye-movements and heartbeats were removed. Followingly, the EEG data were recomposed with the Inverse ICA technique. As a fourth step, the EEG data was re-referenced to the average activity of the two mastoid electrodes (i.e., TP9 and TP10), and the implicit reference was reused as channel FCz. Then, separately for standards and deviants, we segmented the continuous EEG into epochs synchronized to the onset of stimuli from -100 ms before onset to 700 ms after onset. In order to remove remaining artifacts from the EEG data, we applied the automatic artifact rejection algorithm that is included in the BrainVision Analyzer software. During the artifact rejection, the algorithm rejected those segments in which the activity exceeded ± 100 μVs. In addition to this, we calculated the kept and removed segments after the artifact rejection; if a participant’s number of artifacts exceeded 30% of the epochs in each of the conditions, their data was not used in the statistical analysis. After that, the EEG data was baseline corrected (-100 – 0 ms) and finally, the remaining epochs were averaged. Bad electrodes were interpolated with topographic spherical splines. Finally, for the purpose of the statistical analysis, electrode Fz was used as a reference since the specific signal was observed to have been the strongest in most participants and used in previous studies (Pakarinen et al., 2009).

### EEG & Statistical Analyses

First, grand averages of the ERP waveforms were calculated. Figure 1 shows that the deviant stimuli elicited different ERP waveforms at different time-locked events. Furthermore, the difference curves of the ERP waveforms were calculated by subtracting the ERPs of the standard condition from the ERPs of the deviant conditions (see Figure 4).

**Figure 1.**
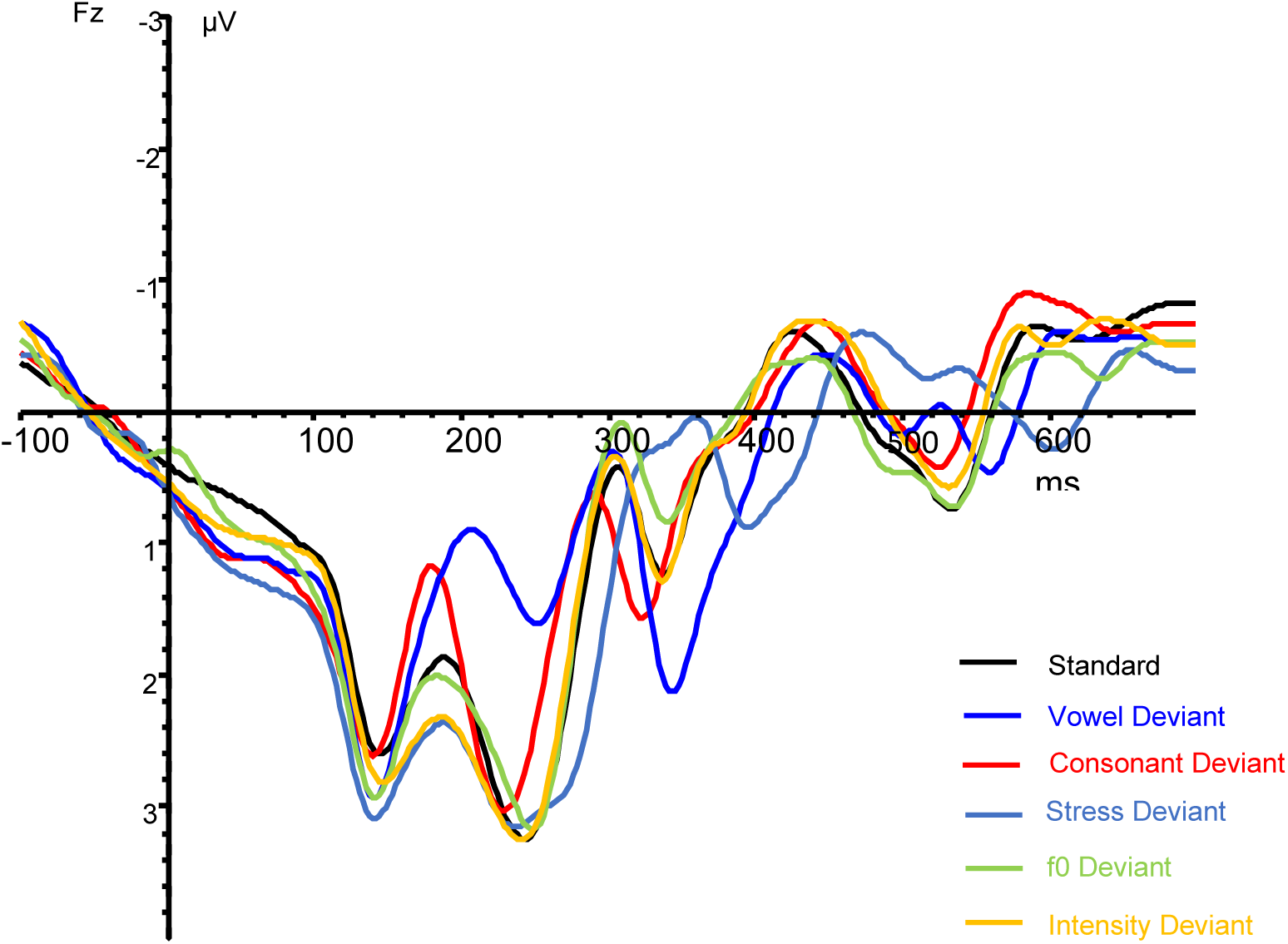
Grand average ERP responses for all stimulus types (standard, vowel, consonant, stress, frequency, intensity) at the Fz electrode. Here and in the following figures, negativity is plotted upwards.

In order to investigate whether MMN peaks were elicited in a statistically significant manner, we performed one-sample t-tests in a range of 176 and 500 ms from stimulus onset on the Fz electrode. With this statistical analysis we wished to examine whether mean amplitudes were different from zero in the aforementioned specified time windows for each condition. For the purpose of the analysis, we selected time windows which showed significant ERP waveforms according to the one-sample t-tests in a timely manner either with eliciting one or two peaks: vowel deviant condition (205 – 255 ms), consonant deviant condition (150 – 200 ms and 235 – 285 ms), stress deviant condition (310 – 360 ms, 510 – 560 ms), F0 deviant condition (205 – 255 ms and 305 – 355 ms), and intensity deviant condition (250 – 300 ms and 445 – 495 ms) (see Table 2). Apart from this analysis, we performed a repeated measures ANOVA to compare the different amplitudes of the MMN ERP components elicited by the deviant conditions. For the purpose of the analysis, we selected only the early time windows (see Table 2) during which the deviant conditions elicited the MMN ERP component. We made this selection since the early time windows allowed the timely natural elicitation of the MMN ERP component. The repeated measures ANOVA was calculated with the factor of Condition (vowel, consonant, stress, frequency, intensity). We used the Greenhouse-Geisser method (G-G) in order to correct the violation of sphericity assumption (Greenhouse & Geisser, 1959). Finally, we used the Bonferroni and Holm tests for post-hoc comparisons and control for the Type I error rate.

**Table 2.** Different time windows for difference curves (i.e., deviant minus standard) for the vowel, consonant, stress, frequency and intensity deviants.

| Deviant Conditions | Peak 1 | Peak 2 |
| --- | --- | --- |
| **Vowel deviant** | 205-255ms |  |
| **Consonant deviant** | 150 – 200ms | 235 – 285ms |
| **Stress deviant** | 310 – 360ms | 510 – 560ms |
| **F0 deviant** | 205 – 255ms | 305 – 355ms |
| **Intensity deviant** | 250 – 300ms | 445 – 495ms |

The one-sample t-tests as well as the repeated measures ANOVA were performed using RStudio and JASP (Version 0.17.1.0).

## Results

First, we checked for normality of the data by conducting the Shapiro-Wilk test; the data were normally distributed. To visualize individual MMN data, we created a series of raincloud plots for each condition. Raincloud plots provide a combination of individual data points along with a boxplot as well as a one-sided violin plot; therefore, they provide transparent statistical information with no redundancy (Allen et al., 2021). In Figure 2, multiple raincloud plots are displayed side by side showing the distribution of the raw data (i.e., the amplitude values) for each deviant condition (i.e., vowel, consonant, pitch, stress, and intensity) in the given time windows.

**Figure 2.**
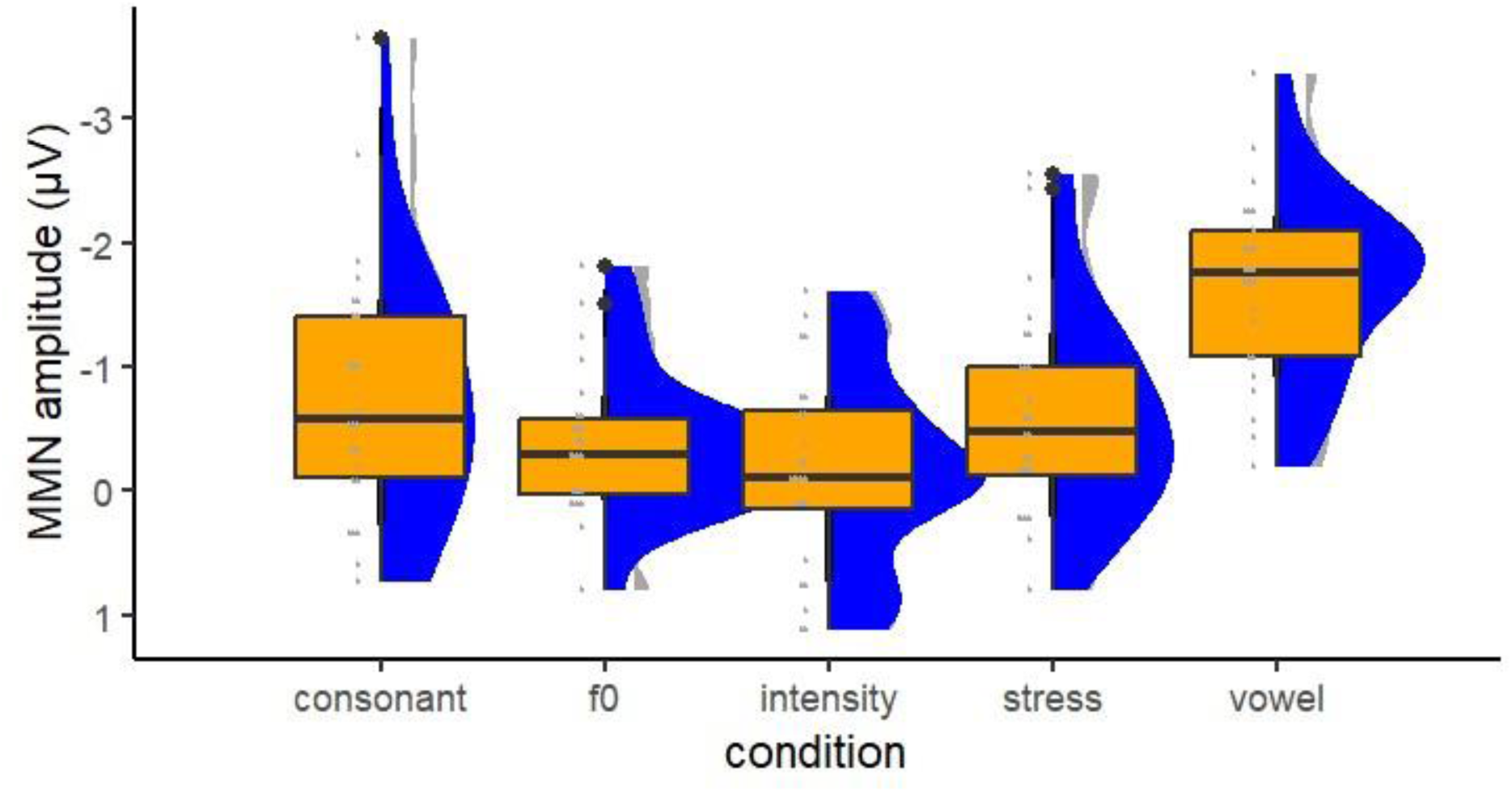
Raincloud plot of the amplitude values of deviants. Each deviant condition is represented with a raincloud or a set of rainclouds if there are multiple selected time windows.

Difference curves of the ERP waveforms were calculated by subtracting the ERPs of the standard condition from the ERPs of the deviant condition. The MMNs peaked between 176 and 500 ms from the stimulus onset. In general, the vowel, the consonant, the stress, the pitch and intensity deviant conditions elicited significant MMN peaks (Figure 4).

Specifically, the vowel condition elicited a statistically significant MMN peak at the time window of 205 – 255 ms (*t*24 = -11.16, *p*<.001, *d* = -2.23). The consonant stimulus elicited statistically significant MMN peaks in early time window of 150 – 200 ms and late time window of 235 – 285 ms (early *t*24 = -3.28, *p* = .003, *d* = -0.66; late *t*24 = -3.80, *p* = .001, *d* = -0.76). The stress condition elicited statistically significant MMN peaks but at later time periods: at 310 – 360 ms (*t*24 = -3.64, *p* = .001, *d* = -0.73) and at 510 – 560 ms (t24 = -5.81, *p* < .001, *d* = -1.17). Pitch and intensity did not elicit statistically significant MMN peaks at the early time windows of 205 – 255 ms (*t*24 = -.92, *p* = .366, *d* = -0.18) and 250 – 300 ms (*t*24 = -1.02, *p* = .318, *d* = -0.20) respectively. Finally, both conditions elicited statistically significant MMN peaks at later time windows; pitch at 305-355 ms (*t*24 = -3.22, *p* = .004, *d* = -0.64) and intensity at 445 – 495 ms (*t*24 = -3.28, *p*=.003, *d* = -0.66).

Repeated measures ANOVA with the factor of Condition resulted in a statistically significant Stimulus main effect, *F* (4, 96) = 20.748, p < .001, η^2^ = 0.464. Mauchly’s test indicated that the assumption of sphericity has been violated, *x^2^* (9) = 13.47, *p* = 0.14, therefore the degrees of freedom were corrected using Huynh – Feldt estimates of sphericity (ε = 0.95).

The raincloud plots in Figure 3 show the significant interactions among the five conditions (i.e., vowel, consonant, stress, f0, and intensity). After the calculation of post-hoc comparisons with the factor condition, the Bonferroni and Holm tests revealed that the vowel was significantly different compared to the other conditions (p < .001) (i.e., consonant, stress, f0, and intensity).

**Figure 3.**
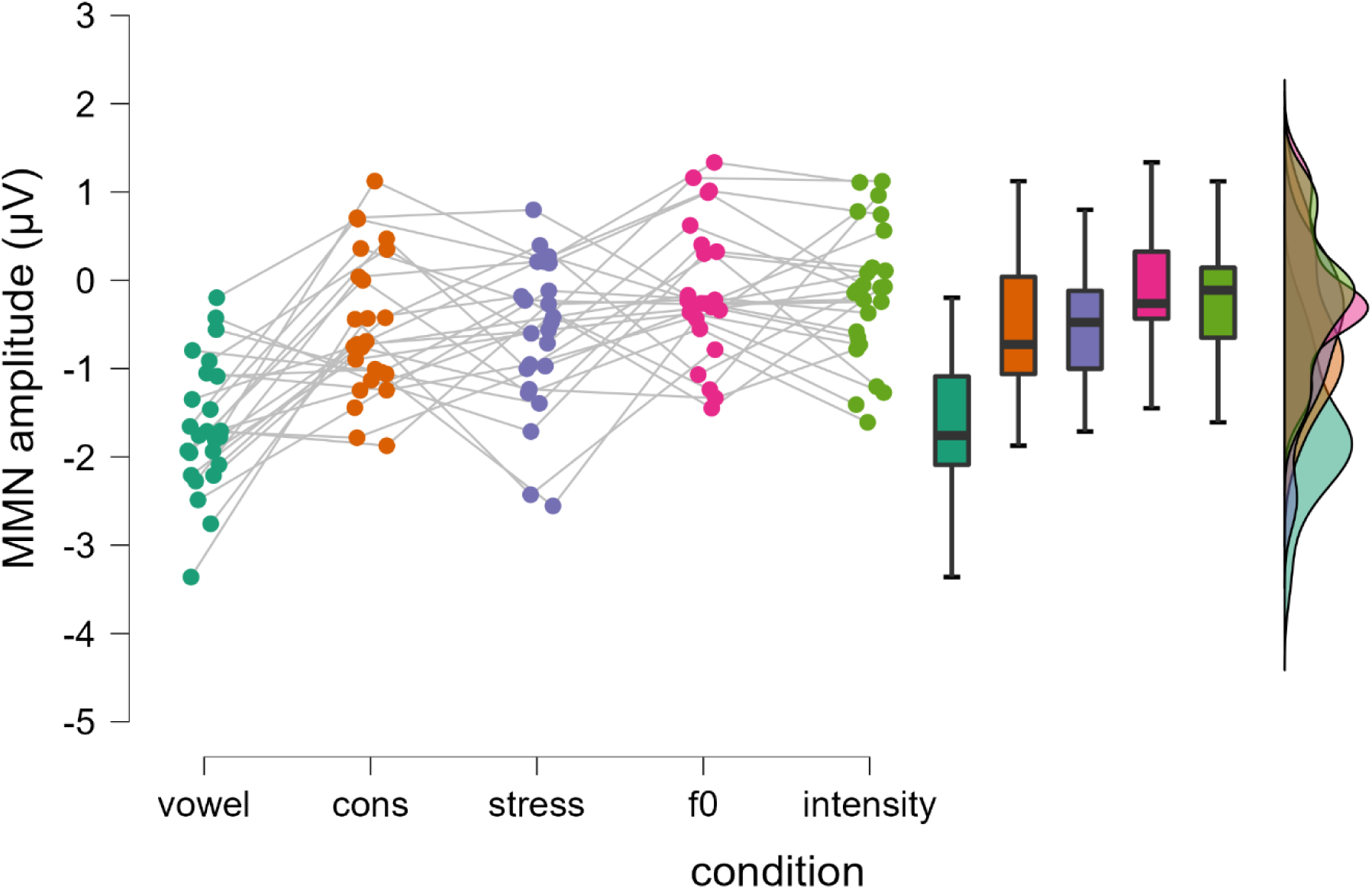
Raincloud plots show the distribution of amplitude among the deviant conditions. The five deviant conditions (i.e., vowel, consonant, stress, f0, intensity) are represented with rainclouds of different colors.

To qualitatively describe the MMN peaks, we created Current Source Density (CSD) maps (see Figure 4). CSD maps depict the magnitude of the current flow from the brain to the scalp (i.e., source) and vice versa (Kamarajan, Pandey, Chorlian & Porjesz, 2015). In the vowel deviant condition, the MMN ERP component was distributed in the frontocentral area of the brain (see Figure 4). In the consonant deviant condition, the MMN ERP component’s left lateralization was more evident in the frontocentral part of the brain (see Figure 4).

**Figure 4.**
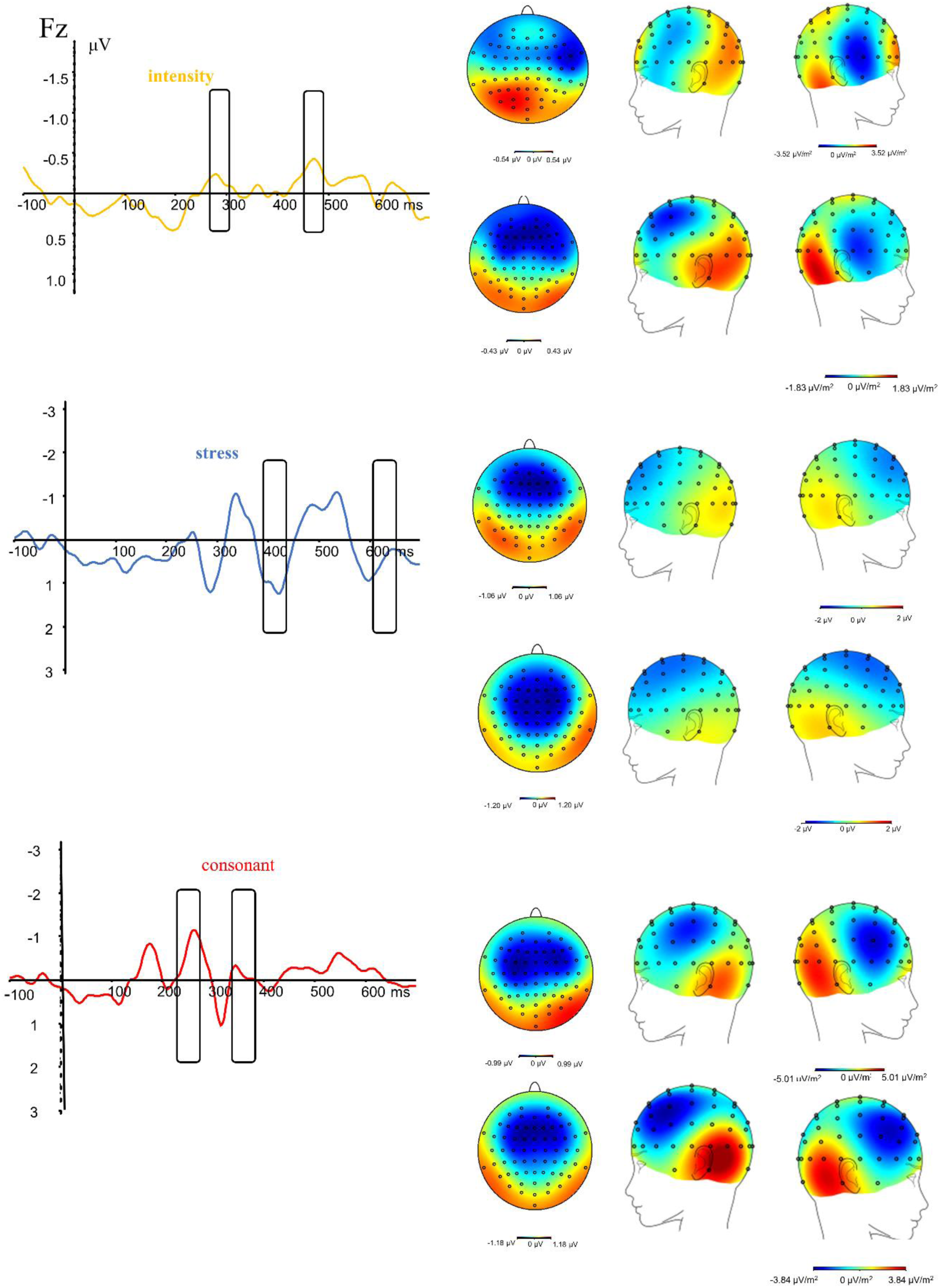
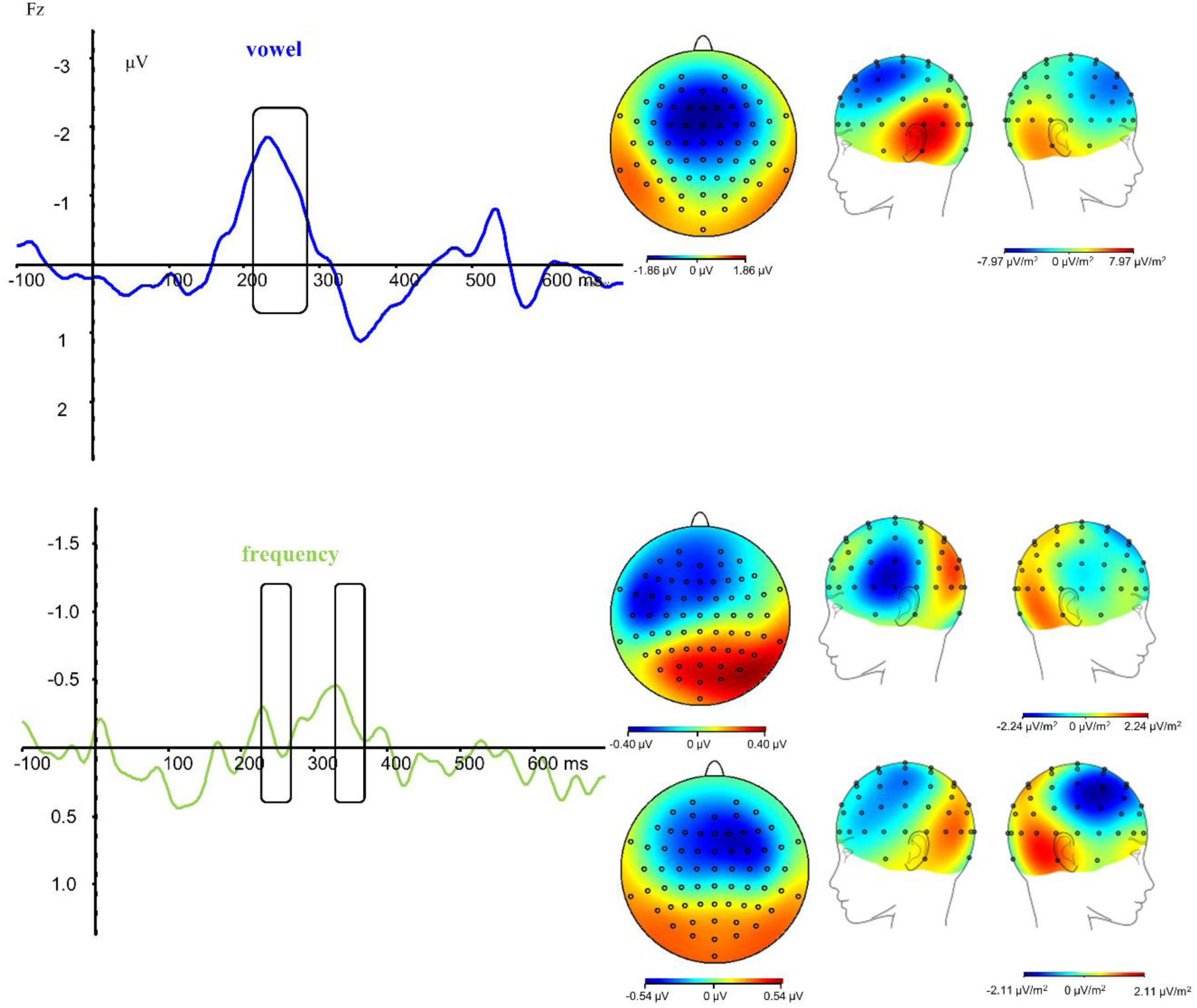
MMN responses are presented for each deviant condition (i.e., vowel, consonant, f0, stress, and intensity). CSD (Current Source Density) maps of MMN ERP component for all deviant conditions, single peak for vowel and two peaks for each other of the deviants (i.e., starting from the top, from left to right: vowel condition, consonant condition, stress condition, f0 condition, and intensity condition). Average activity of the time windows is indicated below the maps.

Frontocentral lateralization was also evident in the stress condition during MMN ERP responses.

## Discussion

The aim of the present study was to develop a multi-feature MMN paradigm which would enable the investigation of the involuntary detection of phonemic, prosodic, and acoustic changes under a short recording time (17 min). All deviant conditions (i.e., vowel, consonant, stress, intensity, and frequency) elicited significant MMN responses with early and late peaks. The signal amplitudes were distributed in the fronto-central brain area focused at the Fz electrode.

The emergence of the MMN ERP peaks after the auditory presence of the vowel and consonant condition appeared statistically significant. This means that the participants were able to passively detect the two phoneme contrasts (/i/ vs. /e/ and /p/ vs. /k/) presented in the disyllabic pseudoword context. Although both the vowel and consonant conditions evoked significant MMN peaks, the vowel condition was the one which showed the highest significance and the largest MMN amplitude (see Fig. 1). This finding is consistent with previous literature (Pakarinen et al., 2009; Sorokin et al. 2010) and it leads to the interpretation that vowel changes may be more discriminable, while other phonological features may be more difficult to detect.

Furthermore, the statistically significant MMN peaks could lead to similar interpretations related to the participants’ performance in the stress deviant condition. The MMN Figure (Fig. 4) demonstrated that two MMN peaks – an early and a later one – were elicited as in the previous conditions. The elicitation of the early MMN peak can -again-explain the participants’ fast ability to detect early changes in prosody. Moreover, the occurrence of the late MMN peak in combination with the earlier one can be interpreted as an LDN ERP peak. In line with previous literature (Honbolygó & Csépe, 2013), the occurrence of LDN in the stress condition might show that stress pattern changes undergo further processing, possibly indicating their interpretation in relation to long-term traces.

Although the vowel, consonant, and stress conditions elicited statistically significant MMN peaks in both early and late time windows, the intensity and frequency conditions elicited statistically significant MMN peaks only in the late time windows but not in the early ones (see Fig. 4). The non statistically significant MMN peaks in the early time windows might give us some insight to Hungarian processing of acoustic elements (i.e., intensity, and frequency). More specifically, this finding might suggest that native Hungarian speakers are not able to detect such changes compared to native speakers of other languages, such as Finnish. Since we followed the exact parameters of the past multi-feature designs customized for the Finnish language (Pakarinen et al., 2009; Sorokin et al., 2010), the intensity and pitch settings of the paradigms may have not been adequately language suited for Hungarian. Further studies are needed in order to clarify why these acoustical changes were not as discriminable by the Hungarian speakers as the rest of the conditions.

Moreover, the CSD maps seen in Figure 4 depict the local electrical brain activity post the MMN ERP component emergence. The left hemispheric positive lateralization in the temporal area for the vowel condition as well as the bilateral one for the consonant and stress condition agree with the previous findings in related studies (Honbolygó et al., 2016). The slight difference in the amplitude distribution of the MMN component indicates a different cortical localization of the MMN for the vowel as compared to the consonant and stress conditions. This suggests that the MMN is not a unitary correlate of speech contrast detection, rather it reflects the acoustic-phonetic differences between the standard and deviant stimuli.

### Limitations & Suggestions

The current MMN multi-feature paradigm showed statistically significant results in most deviant conditions (i.e., vowel, consonant, and stress). However, it did not produce statistically significant results in the remaining conditions (i.e., frequency and intensity). This can lead to the conclusion that the current paradigm might not be equally effective in assessing the phonological processing regarding the parameters set (i.e., vowel, consonant, stress, intensity, frequency). To be more specific, the limitation of the paradigm can be found in its own stimulus design, that of the deviant conditions (i.e., frequency, and intensity). Therefore, a suggestion for improving the paradigm could be to replace these deviant conditions by creating different variations of the vowel and consonant conditions. In this way, the MMN multi-feature paradigm could be more effective in assessing phonological processing in studies with both adults and children.

### Future Directions

The sensitivity of MMN component to different speech features might allow the application of the MMN as a potential biomarker for profiling the central auditory processing of speech sounds. Additionally, the current MMN multi-feature paradigm might be applied as a potential tool for assessing the phonological processing of children and could contribute to their diagnosis with developmental language disorders (e.g., reading disorders or dyslexia).

Following this line, this paradigm can be potentially used in studies with children at risk for reading disorders. Finally, since the improvement in MMN responses may depend on the longevity of experience and training (Bishop, 2007), the present paradigm can be used as an outcome variable to measure the efficacy of applied trainings to children with reading disorders in the context of evidence-based practices (for example, Randomised Control Trial studies).

## Conclusion

In summary, our ERP results demonstrate participants’ involuntary ability to detect linguistic (i.e., vowel and consonant conditions), acoustic (i.e., frequency and intensity conditions), and prosodic (i.e., stress condition) changes. Although our ERP results are consistent with previous literature on the MMN multi-feature paradigm, they were slightly different in the deviant frequency and intensity conditions. This could provide evidence about the processing and discrimination of acoustic changes in native Hungarian speakers compared to native speakers of other languages (e.g., Finnish). Interestingly, the present multi-feature paradigm may lead the way for us to broaden the methods of phonological processing assessment and diagnosis of developmental language disorders; thus, this paradigm could be applied as a potential diagnostic tool for such disorders.

## Acknowledgements

This project has received funding from the European Union’s Horizon 2020 research and innovation programme under the Marie Skłodowska-Curie Grant agreement No.813546. The research was supported by the Research Programme for Public Education Development, Hungarian Academy of Sciences (SZKF-20/2021; PI: FH), and the János Bolyai Research Scholarship of the Hungarian Academy of Sciences [BO/00771/21] awarded to FH. We wish to thank our then lab assistant Dora Bárány and our participants who took part in our experiment.

